# Compositional architecture: Orthogonal neural codes for task context and spatial memory in prefrontal cortex

**DOI:** 10.1101/2025.02.25.640211

**Authors:** JeongJun Park, Charles D. Holmes, Lawrence H. Snyder

## Abstract

The prefrontal cortex (PFC) is crucial for maintaining working memory across diverse cognitive tasks, yet how it adapts to varying task demands remains unclear. Compositional theories propose that cognitive processes in neural network rely on shared components that can be reused to support different behaviors. However, previous studies have suggested that working memory components are task specific, challenging this framework. Here, we revisit this question using a population-based approach. We recorded neural activity in macaque monkeys performing two spatial working memory tasks with opposing goals: one requiring movement toward previously presented spatial locations (look task) and the other requiring avoidance of those locations (no-look task). Despite differences in task demands, we found that spatial memory representations were largely conserved at the population level, with a common low-dimensional neural subspace encoding memory across both tasks. In parallel, task identity was encoded in an orthogonal subspace, providing a stable and independent representation of contextual information. These results provide neural evidence for a compositional model of working memory, where representational geometry enables the efficient and flexible reuse of mnemonic codes across behavioral contexts while maintaining an independent representation of context.

## Introduction

Working memory, the ability to temporarily store and manipulate information, is central to many cognitive tasks^1^. The prefrontal cortex (PFC) has long been implicated in supporting spatial working memory through sustained activity tuned to specific locations^2–7^ across various tasks^8–10^. However, whether a common neural circuit supports working memory across different task contexts or distinct task-specific circuits are required remains unclear. Each possibility has its own advantages. A shared circuit minimizes the number of neurons required and enhances the generalizability of spatial memory coding across tasks, while multiple specialized circuits allow for task-dependent optimizations within the spatial domain.

Recent theoretical work suggests that artificial neural networks reuse core representational structures that can be flexibly adapted to different tasks^11–13^. However, neural evidence for a compositional model is mixed. Some studies report that a single set of prefrontal neurons encodes information similarly across tasks, supporting a compositional model of flexible memory representations^14–17^. Other studies report that different neurons encode information for different tasks, challenging this framework^18–22^. As an example of the latter, one influential study classified neurons based on binary statistical thresholds and found that different sets of neurons were active in a match-to-sample memory task versus a non-match to sample task^22^. However, using a binary significance criterion to classify neurons may exaggerate the extent to which neurons behave differently in two tasks, and so the issue is worth revisiting.

Another key issue is whether and how task information is encoded alongside spatial information in the PFC. If overlapping neural populations represent both task and spatial information (i.e., mixed selectivity^23,24^) and the two representations are not orthogonal, then the task being performed might influence the encoding of spatial information. This would complicate or even preclude the compositional reuse of that representation. In contrast, if task and spatial information are encoded in orthogonal neural subspaces, memory representations could remain stable across tasks and be more easily utilized in a compositional architecture.

To address these issues, we recorded spiking activity from PFC neurons while monkeys performed two spatial working memory tasks requiring different behavioral responses. We found that a single population of PFC neurons consistently encoded spatial memory within a common neural subspace across both tasks while task identity was stably represented in an orthogonal subspace. These findings suggest that the PFC supports working memory through a compositional coding scheme, where spatial information is encoded in a generalizable format while contextual information is independently maintained without disrupting core memory representations.

## Results

### Spatial working memory tasks with different behavioral goals

To compare neural activity in spatial working memory tasks with different demands, we trained two monkeys on both a “look” (memory-guided saccade) and a “no-look” (non-match to sample) task. The task to be performed on a given trial was conveyed by the color of the central cross and visual target (Figure 1, green for the look task and red for the no-look task). Trials began with the monkey gazing at a central cross for 600 ms (fixation period). Next, a peripheral dot (memory target) appeared briefly at a random position along an invisible circle (stimulus period). The monkey was required to remember the target location for at least 1 second (delay period). In the look task, the central cross was then extinguished and the animal was rewarded for shifting its gaze to the memory target location (Figure 1A). In the no-look task, the memory target(s) reappeared along with a novel target, and the animal was rewarded for looking at the novel target (Figure 1B). Look and no-look tasks were presented in alternating blocks of 64 and 192 trials, respectively. In no-look blocks, a second memory target was presented in two-thirds of trials (see Methods for details). The delay period was adjusted to roughly equalize performance in the two tasks (82.18 ± 6.18% accuracy (mean ± SD) in the look task and 81.70 ± 4.54% in the no-look task across 112 sessions). See Holmes et al. 2022 for additional details^25^.

**Figure 1.**
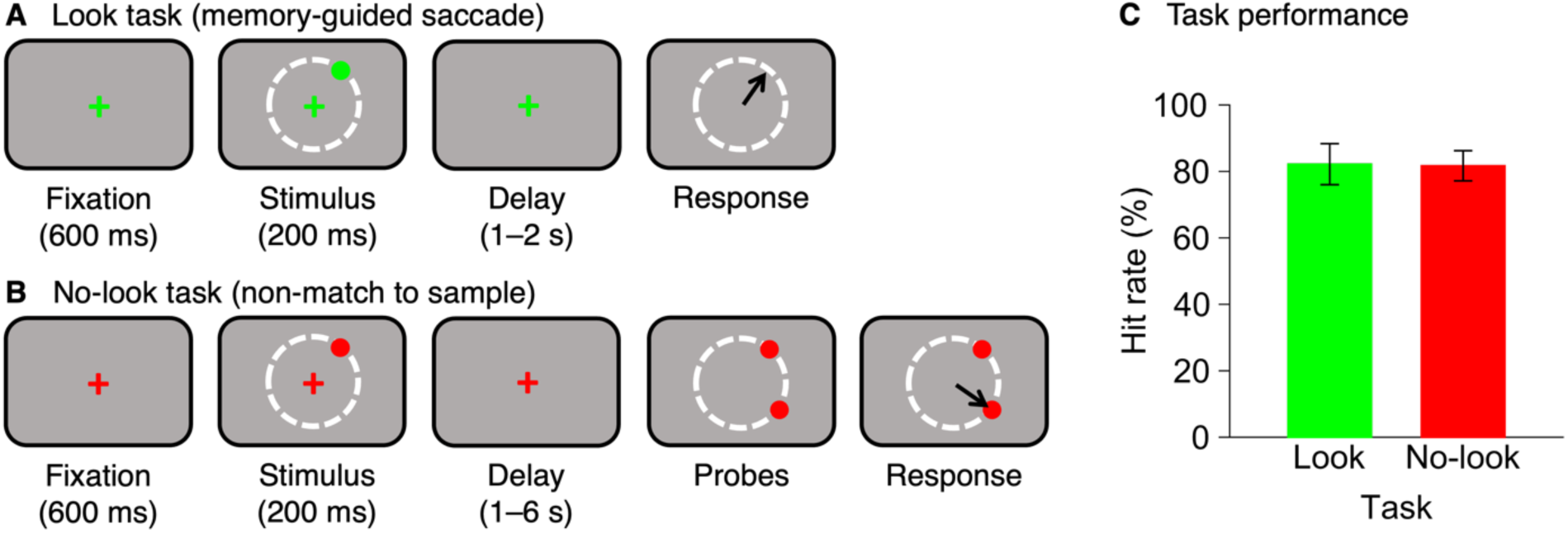
Task design and task performance. (A) Look task: Animals fixate a central green cross. Following a 600 ms fixation period, a green peripheral target appears 13 degrees of visual angle from the central cross in any direction, with a granularity of 1 degree of arc, for 200 ms. After a 1-2 s delay period, the central cross disappears, and animals are rewarded for making a saccade to the target location. (B) No-Look task: Trials are similar to the look task with the following exceptions: all stimuli are red, the memory target reappears along with a novel target at the end of the delay period, and animals are rewarded for making a saccade to the novel target. No-look blocks also included trials with two memory targets (see Methods for details). (C) Mean hit rate in each task. Error bars indicate the standard deviation across 112 sessions.

### Memory tuning in the look task and no-look task

To investigate whether prefrontal cortical neurons encode spatial information consistently across tasks, we compared the memory tuning of individual cells between the look and no-look tasks, focusing on cells with significant spatial tuning during the early delay period (0 to 500 ms from target offset, p < 0.01) in at least one task. We identified 319 memory cells out of 2168 visually responsive PFC neurons from two monkeys and categorized the memory cells as either task-independent or task-dependent, depending on whether they were significantly spatially tuned during the delay period in both tasks or in only one task (Figure 2A). Cells with p < 0.01 in both tasks were labeled as “task-independent” memory cells (27%, n = 86), while those with p < 0.01 in only one task were labeled as “look” or “no-look” memory cells, based on the task in which they showed memory tuning (73%, n = 233; 95 and 138 for look and no-look memory cells). However, these results, like those of the previous study^22^, may overestimate the extent of task specificity. Even if all cells are genuinely tuned in both tasks, less than half (37%) would be classified as task-independent memory cells under a statistical power of 60% and a significance threshold of p < 0.01, whereas if all cells are genuinely tuned in only one task, nearly all (99%) would be identified as task-dependent (Supplementary Table 1). Therefore, relying solely on the statistical significance to assess the task specificity of memory coding is inadequate. Instead, we evaluated the task specificity of memory coding by examining whether the memory tuning properties of all 319 memory cells were consistent across the look and no-look tasks.

**Figure 2.**
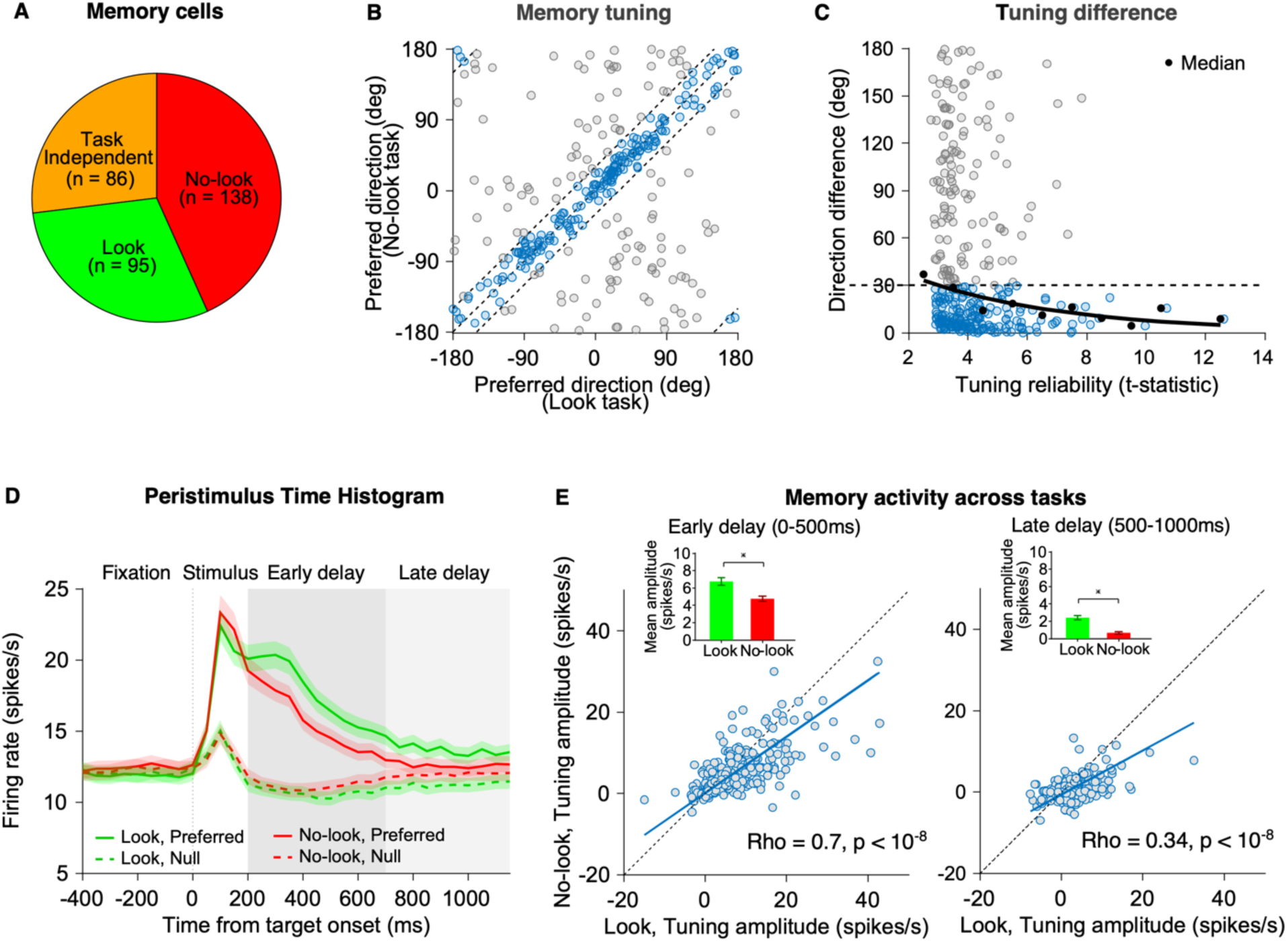
Memory tuning properties in the look and no-look tasks. (A) Categorization of 319 memory cells by their task specificity in memory tuning, based on a significance criterion (p < 0.01). (B) Scatter plot showing the preferred directions of individual memory cells during the early delay period (0 to 500 ms from target offset) in the look and no-look tasks. Cells are color-coded based on the difference in preferred direction across tasks: gray for those with a preferred direction difference of more than 30 degrees and blue for those with a difference of 30 degrees or less. (C) Difference in preferred direction between the look and no-look tasks for memory cells plotted against tuning reliability, measured by uncorrected t-statistics from t-tests comparing firing rates between the preferred and null directions in the task with the larger t-statistic. Black dots represent the median direction differences across all memory cells, binned from 2 to 14 in steps of 1. Black solid line shows an exponential fit to the medians. Cells are color-coded as in (B). (D) Mean firing rate of memory cells, including both task-independent and task-dependent memory cells, over time from target onset in the look task (green) and no-look task (red). Solid/dashed lines indicate firing rates in the preferred/null directions. Shaded regions show the standard error of the mean (SEM). (E) Memory tuning amplitudes during the early (0 to 500 ms from target offset) and late (500 to 1000 ms from target offset) delay periods for individual cells in the look and no-look tasks. Blue solid line shows the Type II regression slope of the scatter plot (early and late delay: 0.69 and 0.54). Inset shows the mean tuning amplitude for each delay period in each task, ±1 SEM.

First, we compared the preferred direction of those memory cells. Most memory cells showed highly consistent preferred directions across tasks, with 59% (188 out of 319) having less than a 30-degree difference in preferred direction between tasks (Figure 2B). This percentage was lower than the chance level of 80% expected if all cells exhibited similar tuning across the two tasks, considering the noise in our measurements (Supplementary Figure 1). However, some discrepancies in preferred direction across tasks may arise due to unreliable estimates of memory tuning. If certain memory cells are genuinely task-dependent, their tuning differences across tasks would persist even with reliable estimates. In contrast, if these discrepancies arise from measurement noise, these differences would decrease as the estimates become more reliable. To test this, we plotted the preferred direction difference between the two tasks against the larger of the two t-statistics for memory tuning (Figure 2C). The result shows that as tuning reliability increases, indicated by higher t-statistics, the discrepancies in preferred direction between the look and no-look tasks decrease. This suggests that the spatial tuning of memory cells is largely conserved across tasks. Thus, while many prefrontal memory cells are categorized as task-dependent based on a significance criterion, the data indicate that most cells are in fact similarly tuned in both tasks.

We next asked whether the amplitude of memory tuning is also similar between the two tasks. We defined tuning amplitude as the firing rate difference when remembering targets in the preferred and null (opposite) directions. Tuning amplitudes were similar in the stimulus period but larger for the look task in the delay period (Figure 2D). Two alternative scenarios that could explain the difference in memory activity between tasks are (1) many memory cells are tuned only in the look task, or (2) most memory cells are tuned in both tasks, but the tuning amplitude is systematically larger in the look task. To address this, we compared tuning amplitudes of individual cells across tasks (Figure 2E). In the first scenario (task-dependent memory cells), we would expect a larger horizontal cluster of look memory cells in the tuning amplitude scatter plot, while the no-look and task-independent cells would form smaller vertical and diagonal clusters, respectively. In the alternate scenario (task-independent memory cells), the tuning amplitudes would form a single diagonal cluster. Our data supported the latter: memory cells formed a single diagonal cluster, with most exhibiting larger tuning amplitudes in the look task (inset in Figure 2E; mean tuning amplitude in the look and no-look tasks, early delay period: 6.75 vs. 4.76 spikes/s, paired t-test, p < 10^−8^, late delay period: 2.41 vs. 0.67 spikes/s, paired t-test, p < 10^−8^). If the amplitude difference reflects a motor preparation signal in the look task that is absent in the no-look task, this effect would be prominent in the FEF, an area closely associated with saccade production. However, this was not the case. In fact, the gain difference was larger in the dlPFC than in the FEF (Type II regression slope of FEF and dlPFC during the early delay: 0.98 versus 0.73, p = 0.025; during the late delay: 0.70 versus 0.55, p = 0.345). Notably, the tuning amplitudes were strongly correlated between the two tasks (Figure 2E; Spearman’s correlation, early delay period: rho = 0.7, p < 10^−8^, late delay period: rho = 0.34, p < 10^−8^). This positive correlation persisted when firing rate was normalized by baseline variability (rho = 0.6, p < 10^−8^ and rho = 0.35, p < 10^−8^, respectively). These results suggest that memory cells collectively represent spatial information with similar weights across both tasks at the population level.

### A common neural subspace for spatial working memory across tasks

We have shown that most cells have similar directional tuning preferences for memory responses in the two tasks, albeit with more robust activity in the look task compared to the no-look task. To directly test the hypothesis that memory representations are similar across the two tasks at the population level, we examined the latent structure of neural population activity underlying spatial memory coding. If the coding axes for spatial memory within the population activity are very similar or identical across the look and no-look tasks, it would suggest that a single neural population code for spatial memory is shared across different tasks. We constructed one dimensional (number of neurons × 1) linear coding axes for spatial memory in each task using the targeted dimensionality reduction (TDR) method^26^ (see Methods). These 1-D coding axes define each cell’s contribution to the neuronal subspaces that represent memory in each task. To quantify whether the subspaces are consistent across tasks, we computed the principal angle between the two coding axes (Figure 3A). Orthogonal axes, that is, axes separated by 90 degrees, indicate independent coding of the two tasks. This could mean that the coding relies on two independent subpopulations of neurons, or that similar neurons are used but that the relative contributions of each neuron are substantially different from one task to the other, or some combination of these two scenarios. Smaller angular separations indicate more similar encodings. Even with identical encoding, sampling error will limit how small the angle between them can be. We estimated the noise floor using a bootstrap on the data within each task. The principal angle across tasks did not significantly differ from the noise floor in either the early delay period (23° vs. 20°, p =0.11) or the late delay period (63° vs. 56°, p = 0.47). These findings indicate that the memory coding axes of PFC population activity were consistent across the look and no-look tasks.

**Figure 3.**
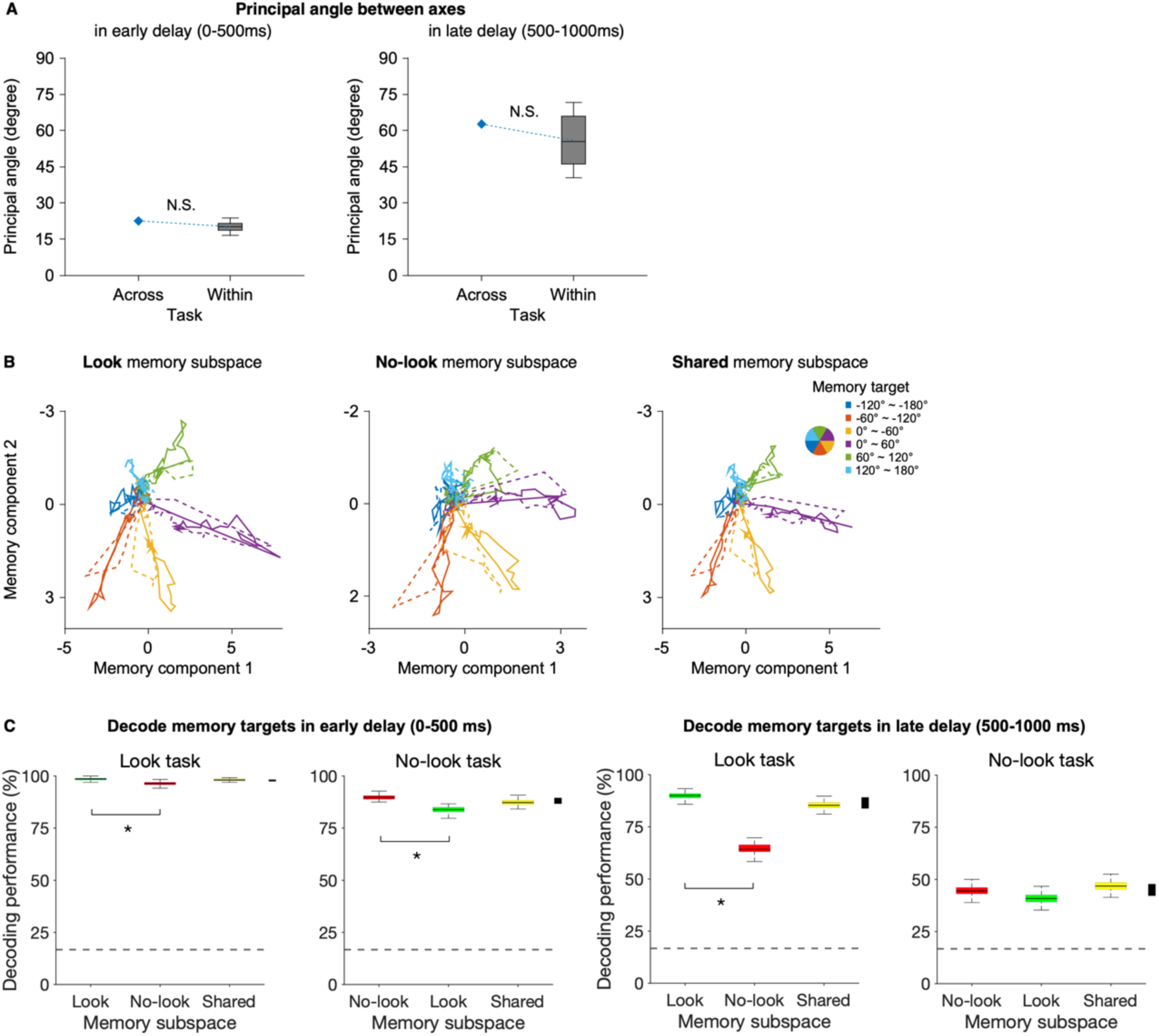
Neural subspaces for spatial working memory between the look and no-look tasks. (A) Blue diamonds represent the principal angle between the memory coding axes in the look and no-look tasks. An angle of 90 degrees indicates independent coding in the two tasks while 0 deg indicates identical coding. Large angles can be caused by noise (sampling error). Gray box plots show the noise floor, obtained by comparing axes within a single task (100 repetitions from each task). In both early and late delay periods, the angle between the look and no-look task axes is not different from the noise floor (p > 0.1). (B) Neural trajectories of population activity of memory cells in the six different target direction bins (see pie chart) from -500 to 1200 ms relative to target onset in the look (left), no-look (middle), and shared memory subspace (right) during the look (solid lines) and no-look (dashed lines) tasks. All three memory subspaces are derived from the delay period activity spanning 0 to 1000 ms after memory target offset. Note the difference in scale for the no-look subspace. (C) Decoding accuracy for target direction in the look and no-look tasks during the early (top; 0 to 500 ms from target offset) and late (bottom; 500 to 1000 ms from target offset) delay periods, using the memory subspace obtained from the look task (green), no-look task (red), or both tasks combined (yellow). Each colored rectangular box represents the interquartile range (IQR) of decoding accuracies across pseudo-trials (see Methods), with the line indicating the median and whiskers extending to 1.5 times the IQR. Dashed lines indicate the chance level (1 out of 6 target direction bins). Black side bars on the far right of each panel show the 95th percentile variation in decoding performance for the subspace trained on the task being decoded (look/no-look memory subspace for the look/no-look task). Differences greater than the side bar heights indicate significant effects (p < 0.05) and are indicated by asterisks.

Thus far, we have quantified mnemonic spatial representations as the difference in firing rate between preferred and null directions, which provides one-dimensional information—either a large or small memory tuning amplitude. However, the actual spatial information was presented in two dimensions on the screen. Therefore, to better capture spatial representation in the PFC, we employed another dimensionality reduction method, demixed principal component analysis (dPCA)^27^, which identifies multiple latent components for task variables while allowing for more than one value per variable. Using this approach, we tested whether 2-D spatial information is encoded in a small number of dimensions within the high-dimensional population response space and whether these dimensions are similar across the look and no-look tasks. We began by splitting our continuous memory targets into six categorical bins, each spanning 60 degrees. We identified latent components of delay period activity that captured the variance in population responses driven by target direction and combined these latent components to define the neural subspace for spatial memory. For visualization, we used the first and second components to construct each 2-D memory subspace. We compared three variants of neural subspaces for spatial memory: one constructed using only trials from the look task (look memory subspace), one constructed using only trials from the no-look task (no-look memory subspace), and one constructed using both tasks (shared memory subspace). If the look and no-look tasks share a neural code for spatial working memory, then the neural representation of target directions should be similar across these three variants. Within each variant (Figure 3B), the trajectories traced out by the two tasks are nearly identical (compare solid to dotted lines). Furthermore, the trajectories are similar across the three variants. In each of subspace, the trajectories associated with contralateral target directions are 2-4 times longer than those associated with ipsilateral target directions. This may, at least in part, reflect the larger number of cells with contralateral mnemonic fields (Supplementary Figure 2). The similar trajectories for the two tasks within and across the three subspace variants support the idea that memory is coded consistently at the population level between the look and no-look tasks.

To quantify coding consistency across tasks, we compared decoding accuracy for memory targets using the three different subspace variants (Figure 3C). We arbitrarily divided the spike data into training and testing sets, then applied dPCA to the training set to separately extract multiple dimensions that capture variation across either time, task or target direction (see Methods). We defined a memory subspace, as above, that would serve as a linear decoder for spatial working memory (see Supplementary Figure 3 for information about the dimensionality of this subspace). We then projected the time-averaged delay period activity of the training trials into this memory subspace to form clusters for each target direction bin. Subsequently, a test trial was projected, and its target direction was decoded as the closest cluster in the memory subspace using a Mahalanobis distance metric. Decoding accuracy was generally high, with no significant difference when using the shared subspace compared to the subspace trained on the task being decoded: 98.1 and 98.5% for the look task in the early period (p = 0.3, permutation test) and 85.3 and 89.8% for the late period (p = 0.1); 87.3 and 89.7% for the no-look task in the early period (p = 0.1) and 46.8 and 44.7% for the late period (p = 1). The average change was only a 1.3% increase in accuracy when the decoder was trained on the task being decoded compared to the decoder trained on both tasks. The effect of training on the alternative task was larger: 98.5 and 96.3% for the look task in the early period (p < 0.01) and 89.8 and 64.5% for the late period (p < 0.01); 89.7 and 83.8% for the no-look task in the early period (p < 0.01) and 44.7 and 40.7% for the late period (p = 0.2). However, this drop of 9.4% in decoding accuracy may reflect overfitting, particularly since the data were collected in blocks. Decoding accuracy still far exceeded the chance level of 16.7% (p < 0.01). Thus, these results are consistent with the use of a common neural population code for spatial working memory across look and no-look tasks.

The preceding analyses were based on cell populations spanning PFC regions. We found similar results when considering only the 88 cells recorded from the frontal eye fields (FEF), and when considering only the 121 cells from dorsolateral PFC (dlPFC). FEF and dlPFC were identified based on anatomical landmarks in magnetic resonance images and on electric microstimulation (sites at which saccadic eye movements were evoked with currents less than 50 μA were defined as belong to FEF)^28^.

### Stable neural representation of task context in PFC populations

We have demonstrated that a single population of PFC cells consistently represents spatial working memory across look and no-look tasks. Given that spatial information is used in diametrically opposite ways in these tasks, the task context must also be represented somewhere in the brain. We asked whether the same neurons that encode spatial memory also encode task context. To address this, we first examined baseline activity. During the initial fixation period prior to stimulus onset, the population-averaged firing rate of the memory cells did not differ by task (Figure 2D, mean baseline activity in the look and no-look tasks: 11.9 vs. 12.3 spikes/s, paired t-test, p = 0.16). However, at the individual cell level, some cells exhibited higher baseline activity in either the look task (Figure 4A) or the no-look task (Figure 4B). Collectively, 48% of memory cells (154 out of 319) exhibited significantly different baseline activity between the two tasks (Figure 4C, Wilcoxon signed rank test, p < 0.05). This proportion was much higher than the 5% (16 cells) expected by chance and also much larger than the proportion of significant differences when comparing random halves of trials within each task (look task: 5.3%, no-look task: 5.6%). The distribution of individual cell differences was symmetric about zero, with a similar number of cells having higher firing in one or the other task, consistent with no difference in average baseline activity between the tasks (Figure 2D). Memory cells in both FEF and dlPFC represented task information, with baseline activity significantly modulated by task in 35% and 37% of cells in each region (31 out of 88 FEF cells and 45 out of 121 dlPFC cells, respectively). Interestingly, the task-induced baseline activity modulation was observed not only in memory cells but also in visually responsive PFC neurons without memory activity (Supplementary Figure 4): 43% of cells with spatial tuning only during the stimulus period and 55% of visually responsive but spatially untuned cells showed significant baseline activity modulation by task.

**Figure 4.**
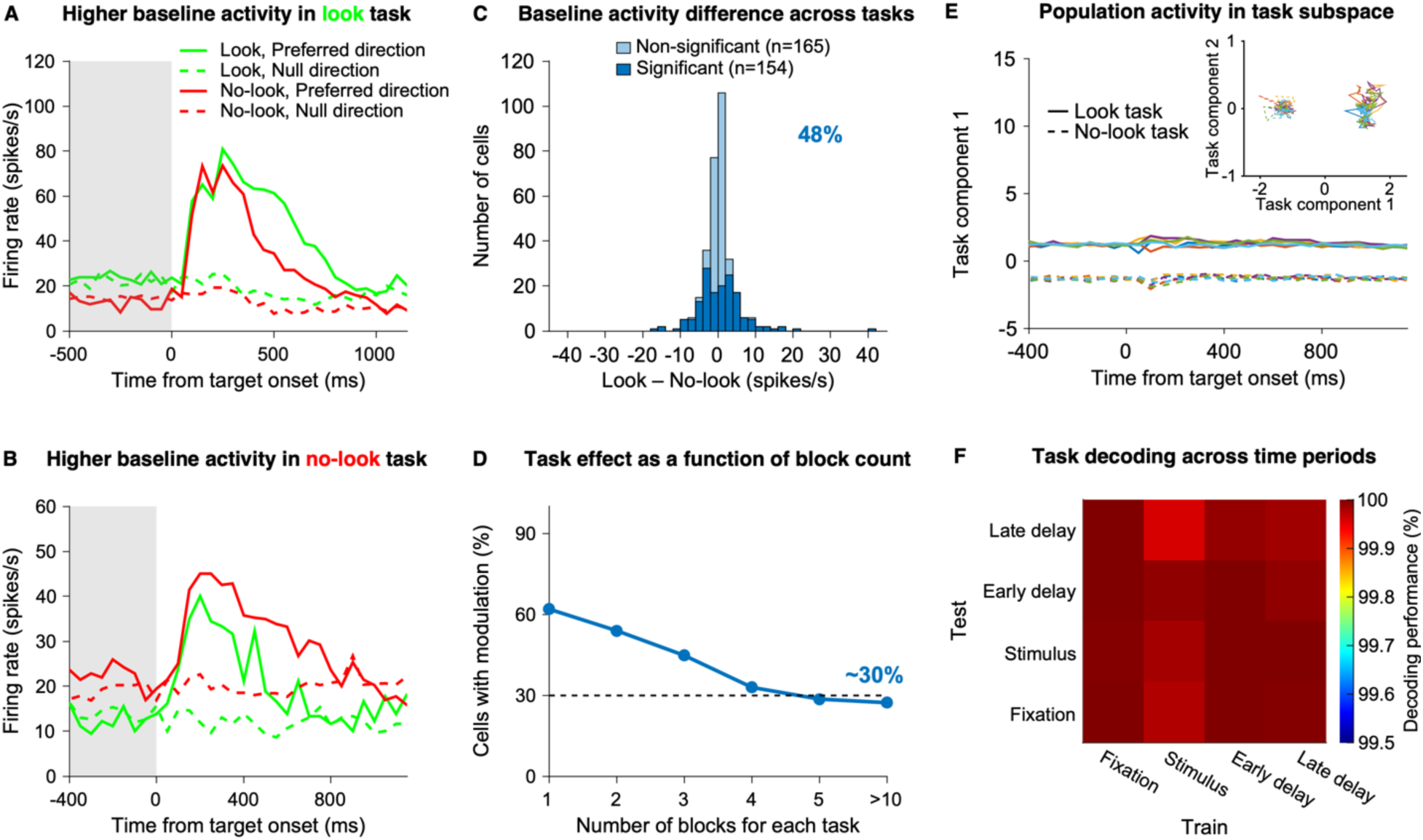
Baseline activity modulation of PFC neurons by task. (A-B) Trial-averaged peristimulus time histograms of example PFC memory cells, showing higher baseline activity during the look task (A) and the no-look task (B). (C) Histogram of the difference in firing rate during the fixation period (−400 to 0 ms from target onset) between the look and no-look tasks (look minus no-look) for 319 memory cells. A total of 154 cells (48%) showed significantly different baseline activity between tasks (Wilcoxon signed rank test, p < 0.05), with similar numbers exhibiting higher baseline activity in one task or the other (82 for the look task and 72 for no-look task). (D) Proportion of memory cells with significant baseline activity modulation by task (Wilcoxon signed rank test, p < 0.05) as a function of the number of blocks collected for each task. (E) Projections of population activity onto the first task component from dPCA as a function of time from target onset, and onto the two-dimensional neural subspace spanned by the first two task components from dPCA (inset). Solid lines represent activity in the look task, while dashed lines represent activity in the no-look task. Each color corresponds to one of six different target direction bins, as shown in the pie chart in Figure 3B. (F) Decoding accuracies for the task across four time intervals: fixation (−400 to 0 ms relative to target onset), stimulus (50 to 250 ms relative to target onset), early delay (0 to 500 ms relative to target offset), and late delay (500 to 1000 ms relative to target offset). Task subspaces were estimated separately for each period, and each subspace was used to decode task information across all four periods, resulting in a 4 × 4 matrix of decoding accuracies.

The use of a blocked design introduces temporal correlations into the trial-by-trial data. Correlations violate the assumptions of our statistical tests and erroneously inflate the significance of our comparisons. If there were no effect of task on baseline firing rate—that is, if the significance we report were due entirely to the block design—then the number of cells showing a significant effect would drop as more and smaller blocks were used. While the proportion of memory cells with significant baseline activity modulation decreased as the number of blocks increased, it reached an asymptote at approximately 30% (Figure 4D). To confirm the existence of task effects, we collected an additional 8 sessions using interleaved 16-trial blocks of the look and no-look tasks from one animal (Monkey F) and then to eliminate any effect of the block design on significance, we compared block means rather than individual trials. Out of 157 cells, 35 (22%) showed significant effects of task (look versus no-look, Wilcoxon signed rank test, n = 10, p < 0.05). In these sessions, we presented no-look trials with two memory targets presented either sequentially or simultaneously in different blocks^25^. When comparing baseline firing in these two no-look blocks as a control, we found only 8 (5.1%) cells showed significant differences, exactly as expected by chance. As a further control, comparing trial-averaged baseline activity across blocks from a single task, randomly divided into two groups, resulted in significant differences in only 7 cells (4.4%), once again not significantly different from the chance level of 5% (binomial test, p = 1). These results strongly support the finding that the baseline activity of PFC neurons is differentially modulated by look and no-look tasks.

Task modulation could remain stable over the course of a trial or change when the memory target appeared. To view task representation as a function of time, we projected population activity onto the neural subspace for task defined by dPCA (Figure 4E). Most of the task information (84% of the total task-related variance) was contained in the first task component (Figure 4E, inset). Notably, within this task subspace, the population activity separated by task remained constant over time, unaffected by memory targets, as expected from a simple shift in baseline activity. To further evaluate the consistency of task representation, we constructed task subspaces from different time intervals, including fixation, stimulus, early delay, and late delay, and attempted to decode task identity across these intervals. Decoding accuracies exceeded 99.9% across all time intervals, compared with a chance level of 50% (Figure 4F). This stable representation of task context was observed in not only memory cells (from both FEF and dlPFC) but also in many other visually responsive PFC neurons without memory tuning (Supplementary Figure 4).

### Independent neural codes for task and memory

A significant fraction of PFC cells’ activity was modulated by both task context and spatial memory, termed mixed selectivity^23,24^. To assess whether task context and spatial memory coding are independent of one another, we compared baseline activity and memory activity. We found no correlation between the task-induced modulation of fixation period activity and the memory-related modulation of delay period activity across cells in either the early and late delay periods, indicating that task context coding was independent of spatial memory coding (blue dots in Figure 5A, Spearman’s correlation, early delay: rho = -0.01, p = 0.8, late delay: rho = -0.05, p = 0.4). Results were similar when measuring memory activity using data from a single task (green and red dots in Figure 5A, Spearman’s correlation: early and late delays using the look task, ρ = -0.02 and -0.03, p = 0.7 and 0.6; early and late delays using the no-look task, ρ = - 0.01 and -0.05, p = 0.9 and 0.3). Further, using the TDR method, we defined the task context and spatial memory coding axes from the fixation and delay periods, respectively (see Methods). We then computed the principal angle between the two coding axes to assess their orthogonality. The angle was close to 90, consistent with orthogonal coding of task context and spatial memory in PFC population activity (Figure 5B).

**Figure 5.**
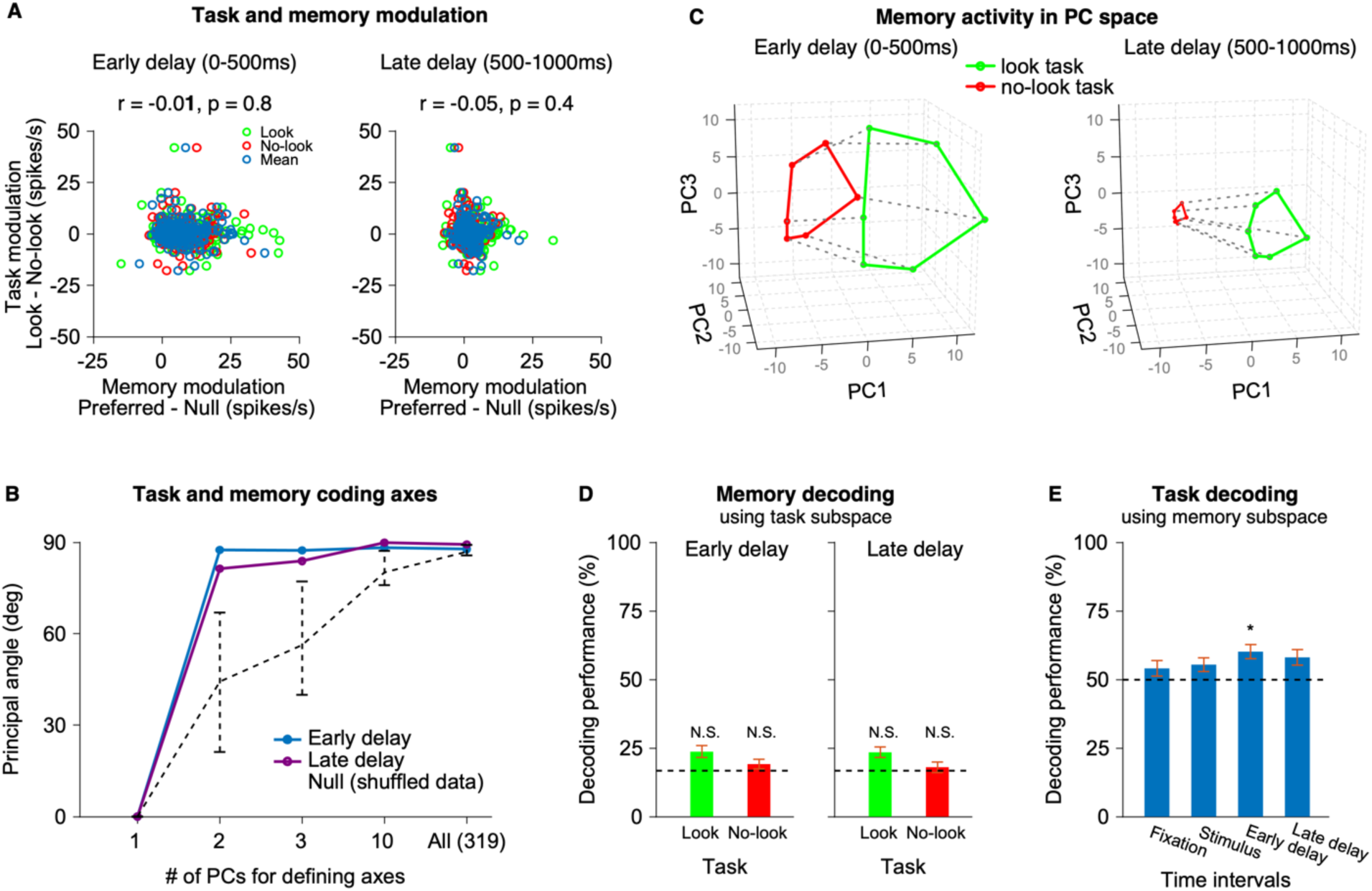
Relationship between task and memory coding in PFC. (A) Scatter plot showing task modulation (baseline activity change, defined as the difference in firing rates between the look and no-look tasks during the fixation period) and memory modulation (memory activity, defined as the difference in firing rates between the preferred and null directions). Memory modulation was computed separately for the look task (green), the no-look task (red), and both tasks combined (blue). (B) Principal angle between the task and memory coding axes, estimated from TDR, as a function of the number of principal components used to define the axes. The blue and purple solid lines represent the principal angle between the task and memory axes during the early and late delay periods, respectively. The black dashed line represents the mean angle between axes from shufled data (1000 repetitions, see Methods). Error bars represent the IQR across 1000 repetitions. The axes are strongly orthogonal when two or more PCs are used. As more PCs are included to define the coding axes, the angle between axes from shufled data increases. Since the first and second PCs almost exclusively contain spatial and task information, respectively, it would be inappropriate to define both the task and memory coding axes using only a single PC (Supplementary Figure 5). (C) Projection of mean firing rates of 319 memory cells during the early and late delay periods into a three-dimensional neural subspace spanned by the first three components from PCA. Each dot represents the population response to one of the six target direction bins, with green and red indicating data from the look and no-look tasks, respectively. (D) Decoding accuracy for target direction using the task subspace during the early and late delay periods for the look (green) and no-look (red) tasks (not significantly different from chance, the dashed line). Compare with Figure 3C. (E) Decoding accuracy for task using the memory subspace during four time periods. Only early delay reaches significance (p < 0.05, asterisk) and the effect size is small (60% correct). Compare with Figure 4F.

To visualize the relationship between the 1-D stable task representation and the 2-D spatial memory representation, we defined a three-dimensional neural subspace spanned by the first three principal components from PCA, using delay period activity (0 to 1000 ms from target offset) from both the look and no-look tasks, and then projected early and late delay period activity from the two tasks onto the subspace (Figure 5C). The locations of the memory targets in the neural subspace reflect an approximate affine transformation of the geometry of the points in real space, that is, a circle. The green circle representing the look task is larger than the red circle representing the no-look task. This corresponds to the difference in memory activity gain for the two tasks (Figure 2E). The two circles are displaced from one another along a nearly orthogonal axis. Based on the simulation models, we found the displacement between two circles corresponds to the task representation, while the size of the circle reflects memory tuning amplitude (Supplementary Figure 6). While the circles corresponding to memory activity shrink in the late delay period, the displacement corresponding to task remains consistent between the early and late periods, reflecting the stable task representation shown in Figure 4E. This result underscores that task and memory information are orthogonally encoded during the delay period.

Finally, as final test of the independence between task and memory coding, we attempted to decode memory information from a task subspace, and task information from a memory subspace. We used dPCA to define the task and memory subspaces. Critically, unlike principal components from PCA, the dPCA components for different variables are not orthogonal by construction, so if task and memory information are not completely independent, information about task will leak into the memory space and vice versa. We attempted to decode remembered target direction by projecting the delay period activity in each of the 6 target direction bins onto the task subspace. The target directions were not decodable in the task subspace during either the early and late delay periods (Figure 5D): decoding accuracy was not significantly different from the chance level of 16.7% (early delay: 24% and 18% in the look and no-look tasks, permutation test, p = 0.1 and 0.2; late delay: 23% and 18% in the look and no-look tasks, permutation test, p = 0.2 and 0.4). Next, we attempted to decode task identity by projected activity from different time intervals, including fixation, stimulus, early delay, and late delay, onto the memory subspace. Once again, the decoding accuracy was not significantly from the chance level of 50%, except during the early delay period, an effect likely due to stronger memory activity in the look compared to no-look task (Figure 5E, permutation test, p < 0.05).

These results support the idea that the neural subspaces for spatial memory and task context are orthogonal.

## Discussion

An efficient way to perform complex tasks is to break them up into smaller pieces and then chain those pieces together in the appropriate way. A compositional model proposes that cognitive processes in neural network rely on a set of shared, combinable components to support a variety of behaviors ^11,12^. While there is strong evidence supporting such a model ^13,14^, there is also strong evidence against the model: a study from Hasegawa and colleagues found that separate PFC circuits exist for storing memorized spatial locations depending on how that memorized information will be used^22^. We posited that the use of significance criteria to classify cells might have distorted their findings and therefore revisited the study using a population-based analysis approach.

After Hasegawa and colleagues^22^, we trained two macaque monkeys on two spatial working memory tasks with different behavioral demands (Figure 1). When we evaluated task participation using a significance criterion, we found that only one-quarter of memory neurons are task-independent, replicating the original study’s results despite minor differences in our methodology (Figure 2A). However, using significance criteria introduces a strong bias to categorize neurons as task-specific (see Supplementary Table 1 for a full explanation). An alternative approach is to ask if cells behave similarly in the two tasks. We find that 59% have similar directional tuning for memory and 41% are mismatched (Figure 2B). However, our method may misclassify cells as mismatched if there is noise in the estimation of tuning directions. Mismatched cells that are very reliably tuned in one task and unreliably tuned in the other would provide strong evidence for task specificity, but we find few of these cells. Most cells with highly reliable tuning in at least one of the two tasks show matched tuning, consistent with task-independent memory cells and a compositional approach to performing complex tasks (Figure 2C). In contrast, most of the mismatched cells do not show highly reliable tuning in either task and so the mismatch may reflect an erroneous estimate of the preferred tuning direction in one or both tasks.

PFC memory cell activity was nonetheless influenced by the task being performed. The amplitude of the memory modulation was twice as large in the look compared to the no-look task (Figure 2D,E). However, this gain effect was similar across cells and had little effect on the population coding axes for spatial information (Figure 3A). A single shared neural subspace encodes spatial memory in both tasks (Figure 3). These results—similar tuning directions in the two tasks, a common population coding axis, and a single shared neural subspace—support the idea of a single memory module being used for our look and no-look tasks, and this in turns supports a compositional approach to performing complex tasks.

Surprisingly, the baseline activity of memory cells was modulated by the task being performed (Figure 4). This modulation was stable throughout the trial (Figure 4E,F) and produced a representation of the task at the population level that was nearly orthogonal to the spatial memory representation (Figure 5). Thus, PFC neurons independently encode spatial memory along with task context.

### Task-independent versus task-dependent coding of working memory

Behavior studies provide evidence for both task-independent and task-dependent memory mechanisms^29–31^. Previous studies argued that separate neural populations in the PFC represent spatial information in different tasks, for example, separate stores for representing a target for an upcoming movement versus a target to be avoided^22^, or for spatial information related to past versus future goals^21^. However, having separate memory stores for every possible task is inefficient, as it requires as many neural modules or subpopulations as the number of tasks to be performed. While distinct mechanisms (e.g., different neural populations or distinct dynamical motifs within the same population^11^) may be necessary for storing different types of information, a single mechanism could suffice for storing similar information across different tasks, even if those tasks ultimately used that information in very different ways. This idea can be generalized to a fully compositional cognitive architecture: rather than wiring an entirely new circuit for each task, existing components can be reused for multiple tasks.

A previous study found evidence against a compositional architecture for spatial working memory: Hasegawa and colleagues used a statistical criterion to classify cells as mnemonically tuned for storing spatial locations in two different tasks^22^. In theory, one can design a paradigm with power approaching 100% (no false negatives) and pair that with a very small significance criterion (no false positives) that would correctly classify neurons. However, because neurons are noisy and animals cannot perform unlimited numbers of trials in a recording session, some cells that are truly tuned in both tasks may be misclassified as tuned in just one (false negative due to insufficient power), while some that are not tuned in either task may be misclassified as tuned in just one (false positives due to a lenient significance criterion). Not only are neurons inherently noisy^32,33^, but isolating multiple single units from each electrical contact is likely to add additional noise^24,34,35^. When we apply a significance criterion, we replicate Hasegawa et al.’s findings—three-quarters of neurons appear task-specific. But when we consider tuning direction or population measures of memory tuning, we find that memory coding is largely task-independent.

An exception to our finding of task independence is that memory activity was stronger in the look task than in the no-look task. This is a form of task-specific coding. Notably, however, the tuning amplitudes of the cells were highly correlated across tasks, suggesting a gain modulation that scales the absolute contributions of individual cells to memory coding while preserving their relative influence. Many readout mechanisms rely on relative rather than absolute influence (as opposed to a threshold mechanism, for example, which may determine whether and when a saccade is triggered^36^), and so a uniform gain is consistent with a single memory mechanism. This interpretation is supported by the fact that a single shared neural subspace captures the population activity encoding spatial memory in both tasks (Figure 3). The difference in gain could reflect the influence of a premotor or movement intention signal that is present in the look but not in the no-look task. Since the FEF is more closely associated with saccade preparation and execution than dlPFC^6,7,28,37^, one might expect a larger difference in tuning amplitude in FEF than dlPFC. However, we found just the opposite. An alternative explanation for the difference in gain could stem from the fact that in the no-look task blocks, a second target would appear in more than half of the trials, while only a single target ever appeared in the look task blocks. Representing two targets at once is known to reduce coding amplitude, likely through a process of mutual inhibition or divisive normalization^38,39^. It is conceivable that a reduction in coding amplitude could occur proactively in our paradigm, such that responses in the no-look task would be reduced compared to the look task even after just a single target is presented. Additionally, future research could explore whether differences in task engagement or related effects such as attention might modulate the amplitude of memory tuning.

### Coding of task context by baseline firing rate

Previous neurophysiological studies have shown that task contexts change the neural activity in the PFC, not only the selectivity for visual objects but also the baseline activity ^40,41^. In this study, we found that the PFC neural population also represents task context through distinct baseline activity patterns during spatial memory tasks (Figure 4). Notably, the representation of task context by PFC baseline activity was independent of PFC’s spatial memory coding: PFC task- and memory-coding axes were nearly orthogonal to one another, and baseline modulation by task occurred in equal numbers in PFC neurons with memory tuning, visual tuning, and no spatial tuning (Figure 5 and Supplemental Figure 4). Critically, the task-induced baseline shifts were balanced across memory cells, ensuring that the population-level mnemonic responses to target directions were identical in the two tasks. For example, a simple vector average of responses would be blind to the shifts in baseline firing of individual neurons, as long as those shifts were independent of preferred direction.

Baseline activity change correlated with task context could assist in cognitive control ^42–45^ in at least one of two ways. First, they might provide a preparatory signal similar to that proposed by Churchland et al.^46^, positioning the entire PFC circuitry at a point within neural space that is appropriate for the particular upcoming task. In this case, for example, the baseline signals could influence how the memory signal is read out, which could in turn support the use of the signal in different contexts. This diverges from a strict version of the compositional model, in which the components for each cognitive variable are independent. A second possibility is that the baseline signals could underlie a representation of the task that would be read out and routed independently from the memory signal and then used to generate task-appropriate responses in a downstream circuit. The idea here would be that two overlapping populations of PFC cells multiplex two independent representations, one coding trial-specific information (spatial location) and the other coding block-specific information (task identity or task rules).

This would be consistent with the compositional model. More work will be required to determine if these signals do in fact contribute to cognitive control, and if so, whether either or both of these mechanisms are involved.

### High versus low dimensional coding of working memory

Neural coding dimensionality (also known as representational dimensionality^24,35^) refers to the number of latent dimensions of neural activity required to represent information. Identifying the dimensionality and geometry of information coding in neural activity is a core challenge for population-level understanding of neural computation^47,48^. Recent studies demonstrated that the neural subspaces for task variables are low-dimensional, including sensory information^26^, working memory^49–51^, and prior expectation^52^. Murray et al.^50^ found that spatial memory was stably represented over time in a low-dimensional subspace within the PFC population response space. During the oculomotor delayed response task, different neural subpopulations represented spatial memory at different time points, reflecting strong temporal dynamics in population activity. Despite these shifts in the active neuron ensembles, the spatial memory representation remained stable in a single neural subspace spanned by the first two principal components of population activity. In line with these results, we found that spatial memory in two very different tasks nonetheless was represented (Figure 3B) and decodable (Figure 3C) in a shared neural subspace spanned by three latent components. Our results suggest that PFC population responses are structured to consistently encode spatial information across tasks (Figure 5C).

The number dimensions required for representing each type of information may increase as the information becomes more enriched. For example, the dimensionality of dlPFC population activity patterns increased with learning more values of objects or actions, indicating that richer information leads to a higher dimensionality of neural codes ^53^. Furthermore, the overall neural coding dimensionality may increase as population activity simultaneously encodes multiple pieces of information^54^. Recent studies^53,55^ pointed out that the representational dimensionality of PFC population codes might be underestimated within a simplified task by ignoring the mixed selectivity^23,24^ of the neurons under complex task conditions. Spaak et al.^55^ found that the dimensionality of PFC population activity was higher during a dual task^56^, which involved simultaneously performing two competing tasks related to spatial attention and memory, compared to performing only one of the tasks.

In fact, many factors may influence our estimate of neural coding dimensionality, including not just task complexity^35,54^, but also methodologic choices such as the spatial-temporal scale of recorded neurons^57–59^ and the choice of dimensionality reduction techniques^26,60^. Stringer et al.^59^ found that coding dimensionality increased with the number of neurons analyzed. Aoi et al.^60^ demonstrated that coding dimensionality depended on the dimensionality reduction method. In our study, we found that the estimate of coding dimensionality depended on the task parameters and dimensionality reduction method used. TDR provided a one-dimensional memory coding subspace, distinguishing preferred and null target directions (Supplementary Figure 5), whereas dPCA (Figure 3 and Figure 5D-E) and PCA (Figure 5C) identified multi-dimensional (at least two) memory coding subspace for two-dimensional spatial locations. Regardless of the coding dimensionality of neural population, the key point is that the spatial working memory is encoded within the same neural subspace across different tasks, consistent with a compositional model of cognition.

## Methods

Two male rhesus monkeys (Macaca mulatta), F and H, participated in the study. All research procedures conformed to the Guide for the Care and Use of Laboratory Animals and were approved by the Washington University Institutional Animal Care and Use Committee.

During the experiments, the animal was seated in a custom-designed primate chair (Crist Instrument, MD, USA) with their head held using a head post (Graymatter Research, MT, USA) in a completely darkened room. Visual stimuli were controlled with customized software and projected by either a cathode ray tube projector (Nippon Electric Company, Japan) or an organic light-emitting-diode television set (LG, South Korea). Eye positions were tracked using an infrared video eye-tracking system (ISCAN, MA, USA).

### Behavioral paradigm

Monkeys were trained on two distinct spatial working memory tasks (Figure 1A,B): a memory-guided saccade task (termed “look task”) and a non-match to sample task (termed “no-look task”). In both tasks, each trial began with the animal fixating a central cross on a screen within 3.5 degrees of visual angle (dva) for at least 600 ms (fixation period). Following the fixation period, a peripheral dot appeared for 200 ms (stimulus period). In early sessions (44 out of 112), the stimulus duration in the look task was 250 ms instead of 200 ms, but we aligned the memory activity data to the target offset to account for this discrepancy. The animal was required to remember the location of the dot, termed the memory target, while it disappeared for 1 to 2 seconds during the look task and 1 to 6 seconds during the no-look task (delay period), while maintaining fixation throughout both the stimulus and delay periods. After the delay period, the fixation cross disappeared, and each task required a different behavioral response to report the animal’s memory (response period). In the look task, the monkeys were rewarded for shifting their gaze toward the memory target location without any guiding stimuli (Figure 1A). In the no-look task, the memory target reappeared along with a novel target, and the animal was rewarded for making a saccade toward the novel target (Figure 1B). A liquid reward was delivered when the animal landed near the correct location within 150 ms of leaving the fixation cross.

Both the memory and novel targets appeared at random locations on an invisible circle with a radius of 13 dva centered on the fixation cross. In both the look and no-look tasks, the location of the memory target was uniformly sampled from 360 points distributed in 1 degree of arc increments. In the no-look task, the novel target appeared at ± 60, ± 120, or 180 degrees from the memory target, resulting in a total of 1,800 (360 × 5) potential target configurations.

We defined successful trials differently for the look and no-look tasks. For the look task, the trial was successful if the animal made a saccade within 12 dva of the memory target location within 150 ms of leaving the fixation cross, at which point the target reappeared and the animal was required to look within 5 dva of the target. For the no-look task, the trial was considered successful if a saccade landed within 4.5 dva of the novel target within 150 ms after leaving the fixation cross.

The look and no-look tasks were alternated in blocks of 64 and 192 trials, respectively. In 128 trials in the no-look task, two memory targets appeared instead of one, either simultaneously, followed by a single delay period (64 trials of two-item simultaneous), or sequentially, with two distinct delay periods (64 trials of two-item sequential). Each task block was differentiated by the color of the fixation cross and the visual targets: green for the look task and red for the no-look task. The number of blocks in a session varied, ranging between 2 and 32. In some early sessions with fewer blocks, blocks could contain up to 500 trials. See Holmes et al. 2022 for additional details^25^. We included the first stimulus and delay periods of the two-item sequential trials for further analyses, as the animals could not differentiate the two trial types (single versus two-item sequential) until the second target appeared. We only used the single memory target data to evaluate the behavioral performance (Figure 1C), though performance on two-item sequential trials was similar^25^.

As a control for the effect of collecting data in large blocks on baseline activity, we collected another dataset (8 sessions) from Monkey F. In each session, the animal performed a total of 30 short blocks, each containing 16 trials: 10 blocks of the look task, 10 of the two-item sequential no-look task, and 10 of the two-item simultaneous no-look task, in an interleaved manner. For the main effect, we compared the block means between the look and two-item sequential no-look tasks, while using the comparison between the two no-look tasks (two-item sequential versus two-item simultaneous) as a control.

### Electrophysiology

In each experimental session, we lowered a single electrode (AlphaOmega, Israel) or a 32-channel vector array (NeuroNexus, MI, USA) into the prefrontal cortex (PFC), including frontal eye fields (FEF) and dorsolateral prefrontal cortex (dlPFC). After allowing a minimum of 30 minutes for the brain tissue to stabilize, we recorded neural activity using the Open Ephys system while the monkeys performed the spatial working memory tasks.

To map anatomical landmarks and distinguish the two subregions, FEF and dlPFC, we used magnetic resonance imaging and electrical microstimulation. Sites where saccadic eye movements occurred in response to currents under 50 µA were classified as part of the FEF. Sites near the boundary between FEF and dlPFC were excluded from subregion classification.

We isolated single units on-line when recording with a single electrode and off-line using an automated spike-sorting algorithm^61^. We then assessed whether each unit responded to visual stimuli by comparing the mean firing rate before (−400 to 0 ms relative to target onset) and after (50 to 250 ms relative to target onset) the memory target appearance. For this and for all subsequent analyses, we first took a square root transform of firing rate to help stabilize variance. To ensure the inclusion of actual cells rather than noise detected by the sorter, we selected units that exhibited a significant response to visual stimuli (paired t-test, p < 0.01). Across 112 recording sessions, we identified 2168 visually responsive cells in the prefrontal cortex of two monkeys: 1451 in Monkey F and 717 in Monkey H.

### Spatial tuning

We characterized the spatial tuning of visually responsive cells during the stimulus (50–250 ms from target onset) and delay periods (0 to 500 ms from target offset) in the look and no-look tasks. We fitted a von Mises function to each cell’s spike responses across target directions to estimate its preferred direction. We then tested whether responses differed significantly between the estimated preferred and null (opposite) directions using a t-test. Running the t-test on the same data used to determine the fit is a type of double-dipping, which, like multiple comparisons, inflates the risk of false positives and requires statistical correction. We determined what this correction should be by repeated testing of randomly shufled data. Under the null hypothesis that there is no difference between the preferred and null directions, a p-value of 0.01 or less should occur in 1% of the shuffles, but we observed ∼4%. Dividing the p-values by a factor of 4 yielded a significance rate close to 1% for shuffled data, and so we adopted this correction factor for the remainder of the analyses. Note that the t-statistics in Figure 2C are uncorrected values.

We identified cells as spatially tuned if the corrected p-value was less than 0.01 for at least one of the two tasks. Based on p-values from the early delay period (0 to 500 ms from target offset), we identified 319 cells with spatial tuning during the delay period (termed *memory tuning*) in at least one task —246 from Monkey F and 73 from Monkey H. Additionally, we identified 362 cells with spatial tuning during the stimulus period (termed *visual tuning*) in at least one task but without memory tuning in either task (303 from Monkey F and 59 from Monkey H). To identify cells without spatial tuning in either task, we selected those with p-values greater than 0.2 during both the stimulus and delay periods in both tasks. This resulted in 180 non-spatially tuned cells, which lacked both memory and visual tuning in both tasks (162 from Monkey F and 18 from Monkey H). Note that the sum of the three groups, 319 + 362 + 180, is less than the total number of units recorded. The remaining 1307 cells had tuning in at least one of the four task/interval combination that was less than p=0.2 but greater than p=0.01.) For analyses that addressed a single task (e.g., Figure 2A-C), we used the preferred direction for that task. For Figures 2D-F and 5A,B, we used the mean of the preferred directions from the two tasks. For analyses that considered activity as a function of time (Figures 2D, 3A-C and 4C), we used time bins of 50 ms.

### Neural population subspace

To identify neural population subspaces related to task context and spatial memory, we began by pooling all memory cells recorded from the two monkeys across different sessions to construct pseudo-trials of population activity. Next, we z-scored the neural activity of each cell and applied three different dimensionality reduction methods to the trial-averaged responses of the pseudo population: principal component analysis (PCA), demixed principal component analysis (dPCA)^27^, and targeted dimensionality reduction (TDR)^26^.

#### Principal component analysis

PCA is an unsupervised dimensionality reduction technique that identifies a linear transformation that maximally concentrates data variance into as few orthonormal components as possible. The first component is the unit vector that maximizes variance when the data are projected onto it. Each successive component is similarly constructed but must also be orthogonal to all prior components. It follows that each component captures less variance than the one before it. By projecting high-dimensional data onto a lower-dimensional space spanned by selected components, PCA can be used to reduce dimensionality while preserving as much variance as possible. For the analysis shown in Figure 5C, we applied PCA to the trial-averaged population activity of neurons across conditions. The conditions included tasks (look and no-look), target directions (preferred and null), and time (50 ms windows spanning −400 to 1200 ms relative to target onset).

#### Demixed principal component analysis (dPCA)

Our goal in using dPCA^27^, a supervised dimensionality reduction technique, is to find latent components in the state space of neural activity that maximize the variance explained by each task-related variable. We identified the components for spatial memory and task context underlying PFC population activity using regression coefficients derived from dPCA. To prevent overfitting while computing the regression coefficients, we applied ridge regression for regularization. Since dPCA accommodates more than two values per variable and provides multiple components for each, we divided all target directions into six bins of 60 degrees, covering the full 360-degree range, rather than limiting our analysis to preferred and null directions. Consequently, latent components for spatial memory (termed *memory components*) and task context (termed *task components*) were derived by applying dPCA to the trial-averaged population activity across the six target direction bins in the look and no-look tasks (neuron × look/no-look task × 6 target direction bins × time). The components for each variable are orthogonal to each other. However, since the components for each variable are estimated independently, the task and memory components are not necessarily orthogonal. For the memory decoding analysis (Figure 3C), we included only the memory variable in dPCA, excluding the task variable, to estimate the memory components (see details in the decoding analysis section).

#### Targeted dimensionality reduction (TDR)

To compare two coding axes—either memory coding axes across tasks, or memory and task coding axes—we computed the principal angle, which quantifies the similarity or alignment between two axes. To compute a single principal angle, we defined the coding axes using another dimensionality reduction method (TDR), which provides a single column vector of denoised regression coefficients (one coefficient for each cell) for a variable of interest.

First, for each cell, within each 50 ms time window, we computed multiple linear regression coefficients for task and target direction using firing rates z-scored across conditions (task × target direction × time). Task and target direction labels were arbitrarily assigned values of -1 and 1, with -1 corresponding to the look task/null direction and 1 to the no-look task/preferred direction. Trials were included only if the target direction was within 30 degrees of the preferred direction (preferred direction trials) or within 60 degrees of the null direction (null direction trials).

We then denoised the regression coefficients by projecting them onto a low-dimensional PC space and then back onto the original N-dimensional neural state space, where N represents the number of cells. The PC space used for denoising could be defined by various sets of PCs, such as a single component (e.g., PC 1) or multiple components (e.g., PC 1 to 4). Including fewer PCs in the denoising space removes more variance (more denoising), whereas including more PCs retains more variance (less denoising).

In Figure 3A, we used PC 1 as it sufficiently captured spatial information (see Supplementary Figure 5). In Figure 5B, different numbers of PCs were used to define the task and memory coding axes: PC1, PC1-2, PC1-3, PC1-10, and all 319 PCs.

We averaged the denoised coefficients for task across the fixation period (−400 to 0 ms from target onset) to define the task context coding axis. Similarly, we defined the spatial memory coding axis by averaging the denoised coefficients for target direction across the delay period: 0 to 500 ms from target offset for the early delay and 500 to 1000 ms from target offset for the late delay.

To estimate the noise floor of the principal angle for the same axes in each task, we randomly split the trials into two halves, defined a coding axis for each set, and computed the principal angle between them. This process was repeated 100 times for each task (Figure 3A, gray bar).

Notably, unlike the original TDR method, we did not apply QR decomposition to forcibly orthogonalize the denoised task and memory coefficients, allowing us to directly measure the degree of orthogonality between coding axes.

### Decoding analysis using subspaces from dPCA

#### Spatial memory decoding

To decode target direction from delay period activity (Figure 3C), we began by splitting the dataset into training and testing sets. For the testing set, we randomly selected one trial per target direction bin for each cell in both the look and no-look tasks, yielding 6 test trials of pseudo-population activity for each task. From the remaining trials, we randomly selected 50 trials per target direction bin for the training set, resulting in a total of 300 training trials (50 trials × 6 direction bins) for each task. Using trial-averaged population activity of the training set, we defined the neural subspace for spatial memory by combining the multiple memory components from dPCA. We then mapped the delay period activity of population in the pseudo-trials from the training set onto the memory subspace, creating 6 distinct clusters for each target direction bin. To decode the target directions of the test trials, we computed the Mahalanobis distances between the test trial activity and each target direction cluster within the memory subspace. Unlike Euclidean distance, Mahalanobis distance accounts for the shape and orientation of the cluster when measuring the distance between a test trial data point and the cluster. The target direction of the closest cluster to the test trial was inferred as the memorized target direction in that trial. We repeated this process 100 times for 60 distinct sets of the 6 target direction bins, with each subsequent set shifted by 1 degree. In total, 36,000 test trials (100 iterations × 60 sets × 6 target direction bins) were used to compute decoding accuracy for each task, with each trial classified as either a success or a failure. In Figure 3C, decoding accuracies were averaged across 60 sets of 6 target direction bins, showing the variance of decoding accuracy across iterations. Note that because pseudo-trials were sampled with replacement, the decoding accuracies are not independent across iterations.

In our memory decoding analysis, the memory subspace functions as a linear decoder for the target direction in memory activity. To compare memory subspaces across tasks, we built separate decoders for each task by applying dPCA to the training dataset of each task and identifying the corresponding memory subspaces. Since we used data from only one task to identify each task-dependent memory subspace, only the memory variable, not the task variable, was included in dPCA, yielding memory components only. We also identified a shared memory subspace by using the combined dataset from both tasks, excluding the task variable to ensure no task information was specified for each trial. Note that the gain modulation effect of tasks on memory activity was taken into account when defining the decoders, as the task-memory interaction component was not removed from the memory component. We used the same number of pseudo-trials for training and testing across all three decoders (look, no-look, shared memory subspaces).

We constructed the task-dependent and shared memory subspaces by concatenating the first three memory components, as the decoding accuracy was saturated even with the addition of more components. The differences in decoding accuracy between task-dependent and shared memory subspaces largely remained consistent as long as at least the first two memory components were included (see Supplementary Figure 3).

To test whether the decoding performance of the other task’s memory subspace and the shared memory subspace differed significantly from that of the memory subspace derived from the task of the test trial, we performed a permutation test. To estimate the chance-level difference in decoding accuracy, we randomly split the trials from the task of the test trial into two sets and separately defined two memory subspaces for each trial set using dPCA. We decoded target directions by projecting the train and test trials to each memory subspace. We randomly split the trials 20 times and averaged the decoding accuracies over all 120 test trials (20 iterations × 6 target direction bins). We repeated this procedure 100 times for 60 sets of the target direction bins, then averaged the decoding accuracies across the direction bin sets, resulting in a 6000-trial null distribution. We calculated the p-value for the difference in decoding accuracy between the memory subspaces for the test trial task and the alternative and shared memory subspaces using this null distribution. We constructed separate null distributions for each delay period (early and late: 0–500 ms and 500–1000 ms from target offset) and for each task (look and no-look).

For memory decoding using the task subspace (Figure 5D), we applied the same method as described above, except the delay period activity was projected onto the task components instead of the memory components. To assess the statistical significance of the memory decoding accuracy of the task subspace, we generated a null distribution of the decoding accuracy by permuting the target direction labels 6,000 times (100 iterations × 60 sets of direction bins). To compute the p-value, we compared the mean decoding accuracy of the task subspace with this null distribution. A separate null distribution was generated for each delay period in each task.

#### Task context decoding

To decode task identity across time intervals (Figure 4F), we used dPCA to estimate task components of the population activity in each interval (fixation: −400–0 ms from target onset, stimulus: 50–250 ms from target onset, early delay: 250–500 ms from target offset, late delay: 500–1000 ms from target offset) using the training set, resulting in four separate sets of task components. The neural subspace for task context (task subspace) consisted only of the first task component, as the second and subsequent components did not contribute to task coding (Figure 4E). We then projected the population activity of the test trials from each of the 4 intervals onto the task subspaces derived from each of the 4 intervals, resulting in 16 combinations. The tasks of the test trials were decoded by calculating the Mahalanobis distance between the test trials and the look/no-look clusters of the training trials in the task subspace, using the same approach as for decoding memory targets. Using 72,000 (100 iterations × 60 sets × 6 target direction bins × 2 tasks) pseudo test trials, we computed the mean task decoding accuracy for each pair of train and test intervals.

## Supplementary Materials

**Supplementary Table 1.**
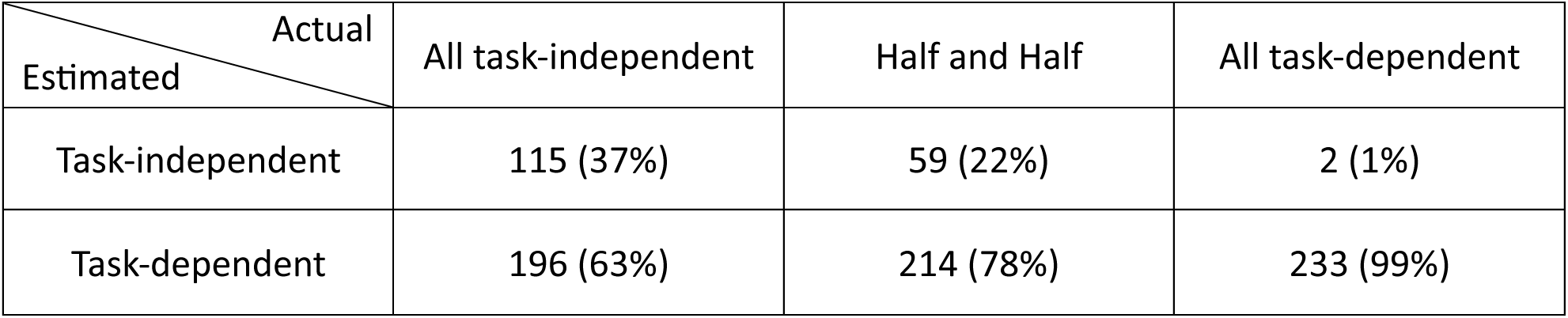
Estimates of task-independent and task-dependent cells. Estimates of task specificity in memory tuning using synthetic data. Following the actual data, the synthetic dataset assumes a total of 2,168 cells, comprising 319 cells with memory tuning and 1,849 cells without memory tuning. Three scenarios for the distribution of the 320 memory-tuned cells were considered:

1) All memory cells are tuned in two tasks (task-independent memory tuning).
2) Half of the memory cells are tuned only in one task, while the other half are tuned in both tasks.
3) All memory cells are tuned only in one task (task-dependent memory tuning).

For each scenario, the number of cells with significant memory tuning in both tasks (task-independent) and in only one task (task-dependent) was estimated, with 60% statistical power and a p-value threshold of 0.01.

Task-independent memory cells in each scenario:

𝐴𝑙𝑙 𝑖𝑛𝑑𝑒𝑝𝑒𝑛𝑑𝑒𝑛𝑡: 115 = 0.6 × 0.6 × 319 + 0.01 × 0.01 × 1849

𝐻𝑎𝑙𝑓 𝑎𝑛𝑑 ℎ𝑎𝑙𝑓: 59 = 0.6 × 0.6 × (319/2) + 0.6 × 0.01 × (319/2) + 0.01 × 0.01 × 1849

𝐴𝑙𝑙 𝑑𝑒𝑝𝑒𝑛𝑑𝑒𝑛𝑡: 2 = 0.6 × 0.01 × 319 + 0.01 × 0.01 × 1849

Task-dependent memory cells in each scenario:

𝐴𝑙𝑙 𝑖𝑛𝑑𝑒𝑝𝑒𝑛𝑑𝑒𝑛𝑡: 196 = 0.6 × 0.4 × 319 × 2 + 0.01 × 1849 × 2 − 0.01 × 0.01 × 1849

𝐻𝑎𝑙𝑓 𝑎𝑛𝑑 ℎ𝑎𝑙𝑓: 214 = 0.6 × 0.99 × (319/2) + 0.4 × 0.6 × (319/2) × 2 + 0.01 × 1849 × 2 − 0.01 × 0.01 × 1849

𝐴𝑙𝑙 𝑑𝑒𝑝𝑒𝑛𝑑𝑒𝑛𝑡: 233 = 0.6 × 0.99 × 319 + 0.01 × 1849 × 2 − 0.01 × 0.01 × 1849

The number of task-dependent memory cells is greatly overestimated, while the number of task-independent memory cells is markedly underestimated. Notably, even when all memory cells are genuinely task-independent, 63% are misclassified as task-dependent, while when they are truly task-dependent, nearly all (99%) are correctly identified as such. Even with 80% statistical power, the analysis is substantially biased, with, for example, percentages of only 59% task-independent and 41% task-dependent cells in the “all task-independent” case.

**Supplementary Figure 1.**
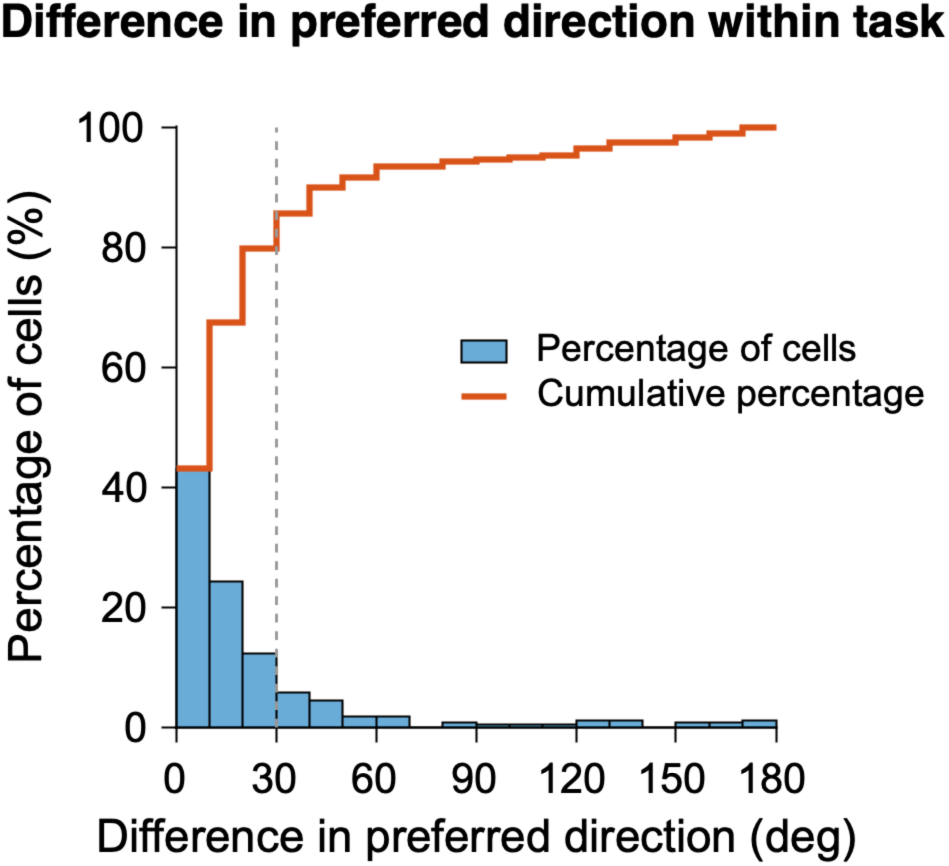
Variation in the preferred direction expected by chance. Difference in the preferred direction of task-independent memory cells between odd and even trials. Cells whose tuning could not be fitted by a von Mises function in either half of the trials were excluded (n = 6 and 4 for the look and no-look tasks, respectively, out of 143 task-independent memory cells), leaving 276 cells (double-counted) in the histogram. The histogram (blue) shows the percentage of cells in each 10-degree bin of preferred direction differences between odd and even trials, while the stair plot (orange) shows the cumulative percentage as a function of the preferred direction difference. 43% of cells have less than a 10-degree difference, 67% have less than a 20-degree difference, and 80% have less than a 30-degree difference.

**Supplementary Figure 2.**
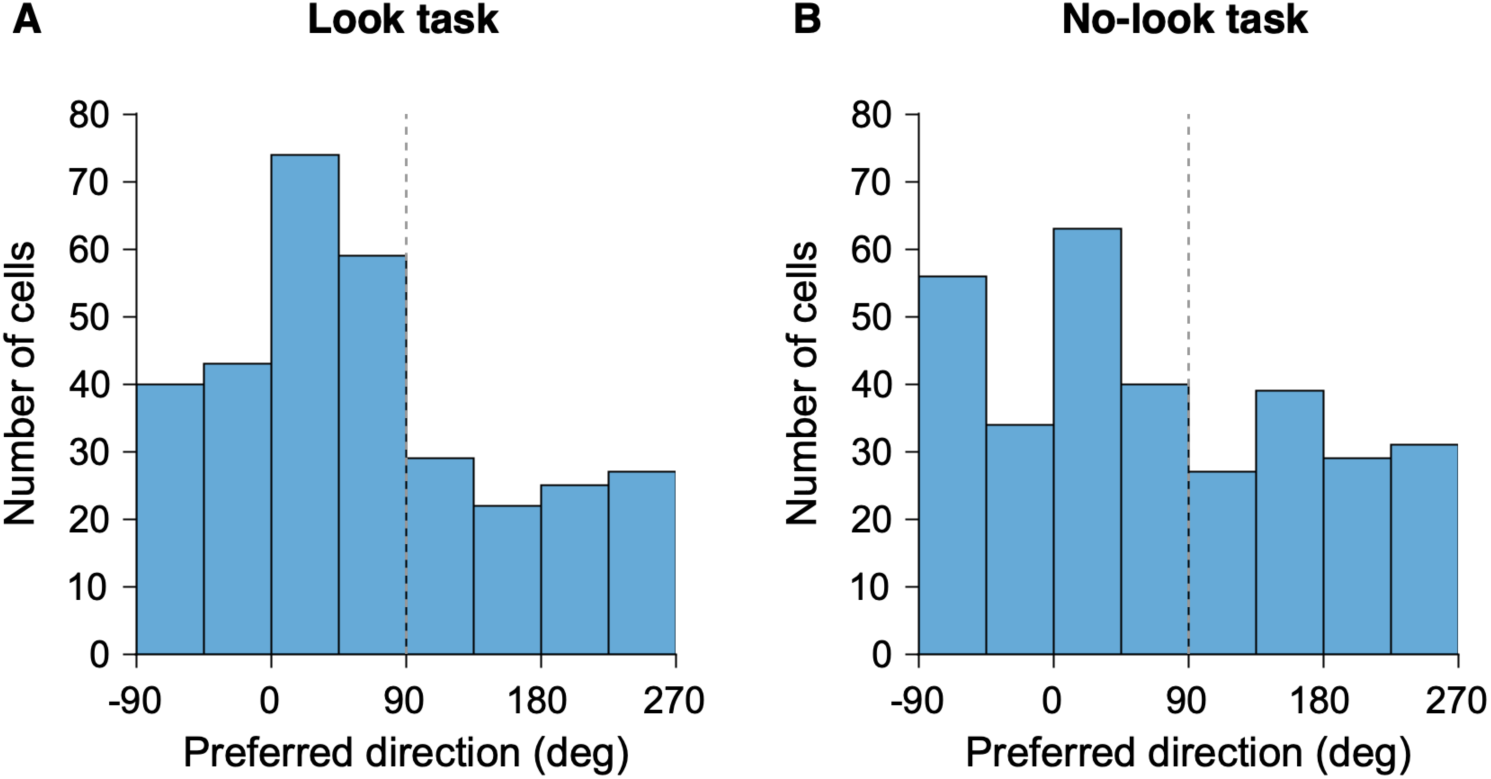
Spatial tuning of memory cells. (A) Histograms showing the preferred target directions, which are the spatial locations of memory targets with the highest firing rates, for 319 memory cells during the delay period (0 to 500 ms from target offset) in the look task. (B) Same as (A), but for the no-look task.

**Supplementary Figure 3.**
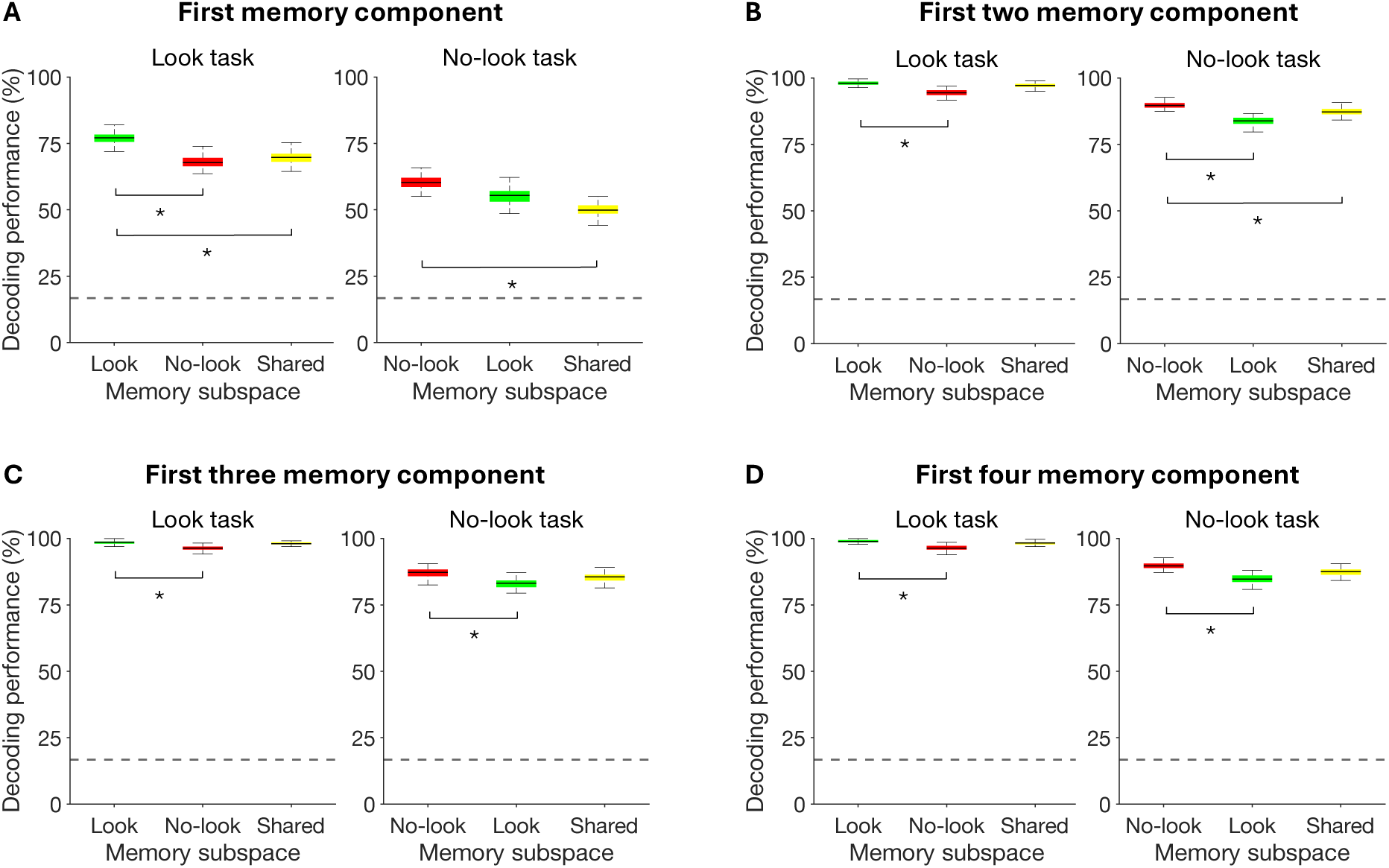
Memory decoding accuracy using different numbers of components. Decoding accuracy for target direction in the look and no-look tasks during the early delay period (0 to 500 ms from target offset), using memory subspaces derived from the look task (green; look memory subspace), no-look task (red; no-look memory subspace), and both tasks combined (yellow; shared memory subspace). Memory subspaces were constructed by combining 1 (A) to 4 (D) memory components from dPCA. Decoding accuracies for all three memory subspaces increased with the inclusion of more components but saturated after the first three components, and the decoding performance of the shared memory subspace was no longer significantly different from that of the memory subspace from the task of the test trial (permutation test, p > 0.1, see Methods for details). Therefore, we used the first three memory components for Figure 3C. Asterisks indicate significant difference between two memory subspaces (permutation test, p < 0.05).

**Supplementary Figure 4.**
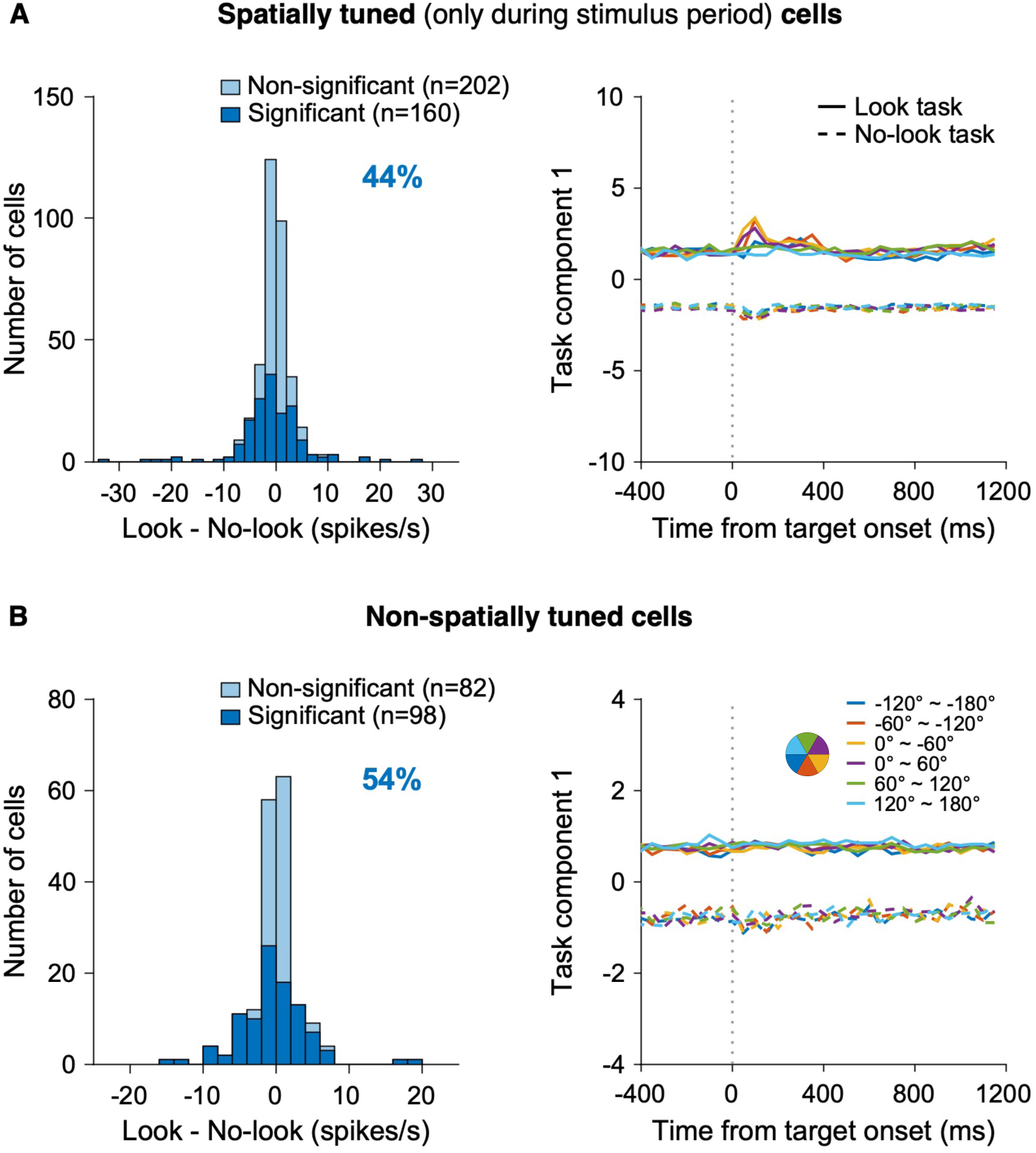
Task-induced baseline activity modulation of other PFC neurons. (A) Left: Histogram of difference in firing rate during fixation period (−400 to 0 ms from target onset) between the look and no-look tasks (look minus no-look) for 362 PFC neurons, which are visually responsive and spatially tuned only during the stimulus period (50 to 250 ms from target onset). 44% (160 out of 362 cells) exhibited significantly different baseline activity between tasks (Wilcoxon signed-rank test, p< 0.05), with 62 cells showing higher activity in the look task and 96 in the no-look task. Right: Projection of population activity onto the first task component from dPCA as a function of time from target onset. Solid lines represent activity in the look task, while dashed lines represent activity in the no-look task. Each color corresponds to one of six target direction bins, as illustrated in the pie chart in (B). (B) Same as (A), but for 180 PFC neurons showing visual responses but not spatial tuning. Left: 54% (98 out of 180 cells) exhibited significantly different baseline activity between tasks (Wilcoxon signed-rank test, p< 0.05), with 43 cells showing higher activity in the look task and 55 in the no-look task. Right: Projection of population activity from 180 non-spatially tuned PFC neurons across six directions in two tasks onto the first task component from dPCA, plotted as a function of time from target onset.

**Supplementary Figure 5.**
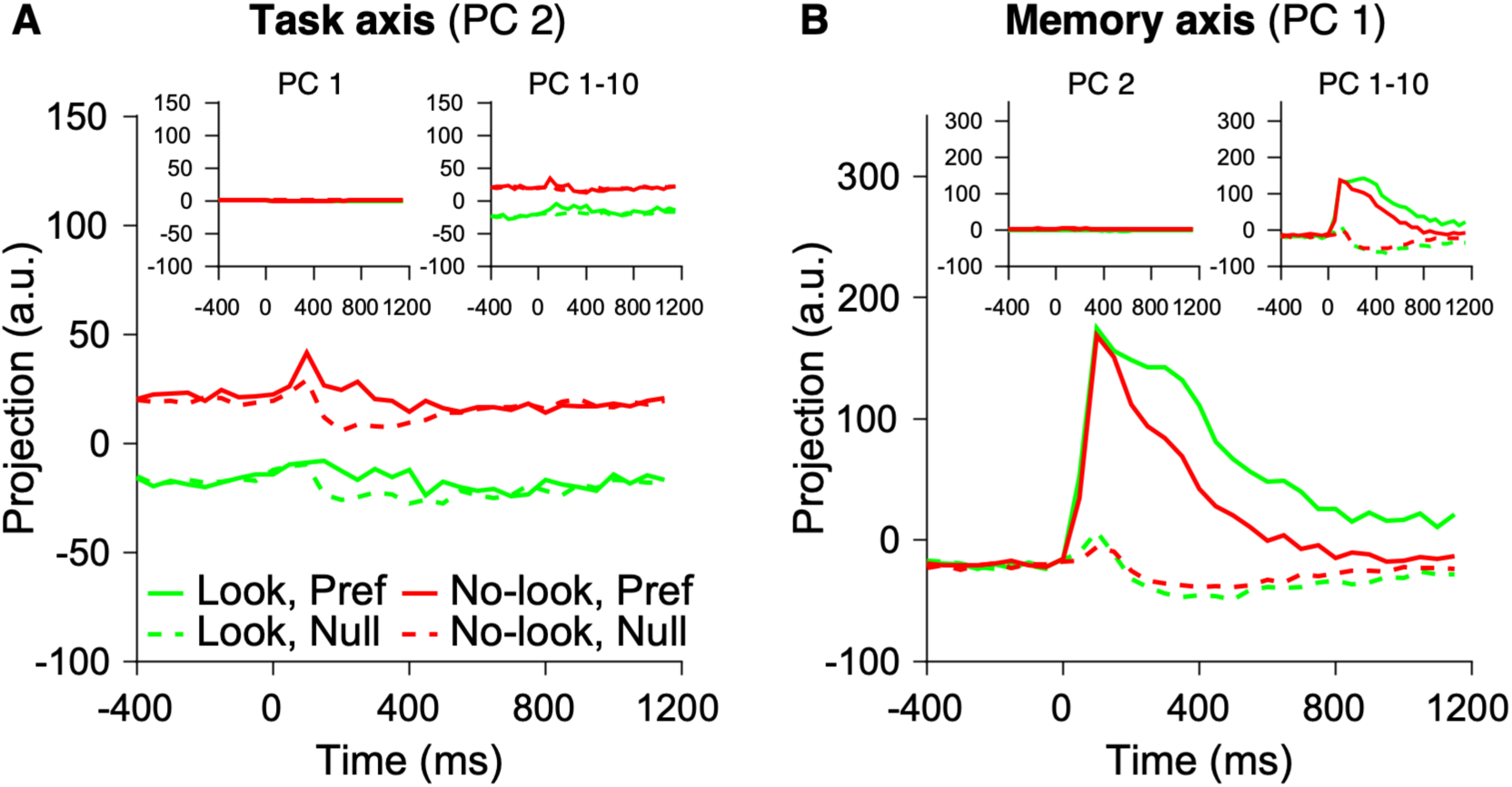
Population activity on the task axis and memory axis. We defined one-dimensional coding axes for task and memory using the targeted dimensionality reduction (TDR) method, where coding axes for each variable are derived from regression coefficients denoised by a subset of principal components from PCA (see Methods for details). (A) Projection of population activity onto the task axis, derived from the TDR method, plotted as a function of time relative to target onset. The task axis was defined using the regression coefficients for task during the fixation period (−400 to 0 ms from the target onset), denoised by the second principal component (PC). Insets show projections onto task axes defined using just PC 1 and the first ten PCs, respectively. Task information was represented on the task axis when the task coefficients were denoised on PC 2, but not when denoised on PC 1. Using the first ten PCs shows a slightly more stable task representation across time and memory targets. (B) Projection of population activity onto the memory axis, defined by the denoised regression coefficients for target direction based on delay period activity (0 to 500 ms from the target offset). Spatial information was well represented on the memory axis when PC 1 was used to denoise the memory coefficients, while adding PC 2 and other components had little impact, indicating that PC 1 captures most of the variance in the memory activity. This suggests that the dimensionality of the memory axis underlying PFC population activity may be 1D, at least for representing the two target directions: preferred and null.

**Supplementary Figure 6.**
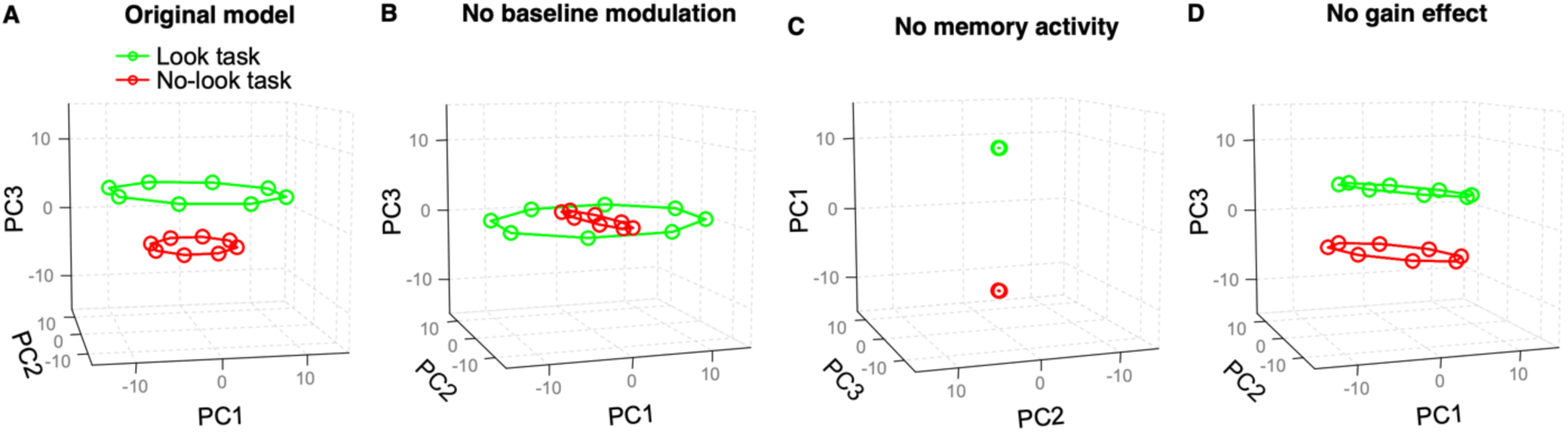
Simulations of neural activity in PC space during the delay period. We modeled the activity of 100 neurons across 100 trials in the look and no-look tasks, respectively, using the equation: 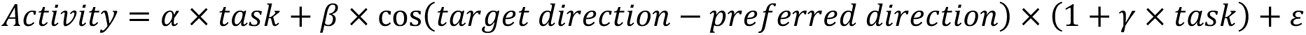

where *task* is categorical variable with values of -1 and 1 for the look and no-look tasks, *target direction* is one of eight directions ranging from 0 to 2 𝜋 radians in 𝜋/4 increments, and *preferred direction* is randomly selected from a uniform distribution between 0 and 2 𝜋 radians. The three parameters (𝛼, 𝛽, 𝛾) represent different neural modulations: 𝛼 indicates baseline activity modulation by task; 𝛼 ∼ 𝑁(0,5), 𝛽 indicates memory tuning strength; 𝛽 ∼ 𝑁(0,20), and 𝛾 indicates the gain of memory activity across tasks; 𝛾 ∼ 𝑁(−0.5,0.15). 𝜀 represents the gaussian noise; 𝜀 ∼ 𝑁(0,0.1). We averaged each model neuron’s activity across trials and z-scored the trial-averaged activity using the mean and standard deviation across all task × target direction conditions. We then applied PCA to the z-scored trial-averaged activity and projected the population activity onto the first three principal components. (A) shows the simulation of neural representation in the 3-D PC space, when the neurons have baseline activity modulation, memory tuning, and gain effects. (B-D) shows the neural representation in the subspace where baseline activity modulation (𝛼 = 0), memory tuning (𝛽 = 0), gain effect (𝛾 = 0) is removed, respectively. (B) Removing baseline activity modulation aligns the two task representations on the same plane while preserving their shape and size. (C) Removing memory tuning disrupts the circular shape of the spatial representation, although the two task representations remain separable. (D) Removing the gain effect makes the spatial representations identical while maintaining their distance from each other.

## Acknowledgments

We thank H. Song for helpful discussion and feedback regarding this work and K. Duckworth and J. Tucker for helping with technical issues and collecting data.

## Author contributions

JP analyzed the data and wrote the manuscript. CH designed, conducted the experiment, and collected the data. LS helped design the experiment and write the manuscript.

## Funding

This work was funded by the National Eye Institute (grant number: R01-EY012135), the National Science Foundation (grant number: T32-NS073547), and the National Institute of Mental Health and the National Institute of Biomedical Imaging and Bioengineering (grant number: R01-EB028154-01).

## Conflict of interest

The authors declare that the research was conducted in the absence of any commercial or financial relationships that could be construed as a potential conflict of interest.

## Notes

### Competing Interest Statement

The authors have declared no competing interest.

